# Building methodological consensus to ensure rigor and reproducibility in zebrafish fertility research

**DOI:** 10.1101/2023.06.19.545565

**Authors:** ME. Kossack, K. Bowie, L. Tian, JS. Plavicki

## Abstract

Zebrafish are an increasingly popular model for studying the genetic and environmental factors that shape male and female fertility; however, the field currently lacks a standardized approach to fertility assessment. The current lack of consensus makes comparisons across studies more challenging and is an obstacle to reproducibility in the fields of reproductive biology and toxicology. Here, we review the diversity of spawning approaches used in zebrafish reproductive toxicology research to asses fertility and provide evidence that spawning parameters can result in meaningful differences in egg production and spawning success.

**Highlights:**

- Zebrafish fertility research lacks methodological consensus
- Measures of fertility vary with age and frequency of spawning
- Methodological consensus will increase reproducibility in toxicology

## Main text

In an attempt to build on previously published results, we replicated an exposure paradigm previously shown to reduce female fertility. However, we found no significant effect of toxicant exposure on female fertility. While we considered that differences in genetic background and environmental differences inherent to using different fish facilities could shape the outcome of our studies, we also asked whether the fundamental parameters in fertility assessment such as the frequency of spawning, individually paired versus grouped spawns, and age of spawning pairs could significantly influence the outcome of fertility assessments. To determine how variable these experimental parameters were across studies, we performed a literature search on PubMed using the terms (“zebrafish” OR “danio rerio”) AND (“fertility” OR “infertility” OR “reproduction” OR “spawn” OR “eggs released”). The search yielded a total of 1545 publications at the time of the search. Studies were excluded if they did not use zebrafish as a model organism, spawn fish, include a method of assessing fertility, use a control group, or if they assessed fertility through histology. Studies that met the previously defined criteria were added to Table 1 in the order of their appearance in PubMed. After evaluating 20 publications, we found that there was a high degree of variability in the methodology used to assess fertility (**Table 1**).

**Table 1.** Prevalent fertility testing conditions in the published literature. Surveying the literature, we found there is no consistency in the methodology used to assess fertility in zebrafish. Abbreviations used in the table: months post fertilization (mpf) and hours post fertilization (hpf).

|  |  | Ref |
| --- | --- | --- |
| **Male to Female Ratio** | **Individually paired spawns** | [1–6] |
|  | **Groups: Equal ratio of males to females**<br>2:2, 3:3, 5:5, 15:15, 2:2, 8:8, 9:9, 4:4, 10:10 | [7–18] |
|  | **Uneven sex ratios**<br>7:5, 2:3 | [19,20] |
| **Frequency** | **Continuous co-housing**<br>7, 14, 21, 10, 30, 36 days | [2,6,8,10–14,17–19] |
|  | **Weekly** | [1,3,4,9] |
|  | **Biweekly** | [7,20] |
|  | **Other** | [5,15,16] |
|  | *Every other day, Once, Inconsistent* |  |
| **Age at first spawning** | **3 mpf** | [16,18,20] |
|  | **4 mpf** | [1,7,13–15] |
|  | **5 mpf** | [3,9,12] |
|  | **6–7 mpf** | [8,10] |
|  | **Other**<br>“>6 mpf”, 12 mpf, “Adult”, 8 mpf, 15 mpf | [2,4–6,11,17,19] |
| **Age of embryo at time of fertilization assessment** | **Within Hours**<br>2 hpf, 3 hpf, 4–6 hpf, 30 mins | [1–3,6,9,10,12,15,17,19] |
|  | **24 hpf** | [4,7,8] |
|  | **“Daily”** | [11,13,14,18] |
|  | **>24 hpf**<br>3 dpf, 5 dpf | [5,16,20] |

We performed a series of experiments to test the hypothesis that spawning frequency alters measures of fertility. Group spawns with equal sex ratios and individually paired spawns are the most frequently used mating paradigms (**Table 1**). We tested the effects of weekly, biweekly, and monthly spawning on individually paired spawns. Fertility was assessed using the number of eggs released and the fertilization rate of the eggs produced. We found that there was no significant effect of age on the number of eggs released (p=0.46301, linear regression); however, there was a significant effect of frequency (**Figure 1A**). These data suggest that studies using the total number of eggs released as a marker for fertility can be pooled across ages, but not all spawning frequencies should be considered equivalent experimental parameters. Weekly individually paired spawns yielded significantly fewer eggs than biweekly or monthly individually paired spawns (p<0.05, one-way ANOVA with Tukey’s post-hoc test).

**Figure 1.**
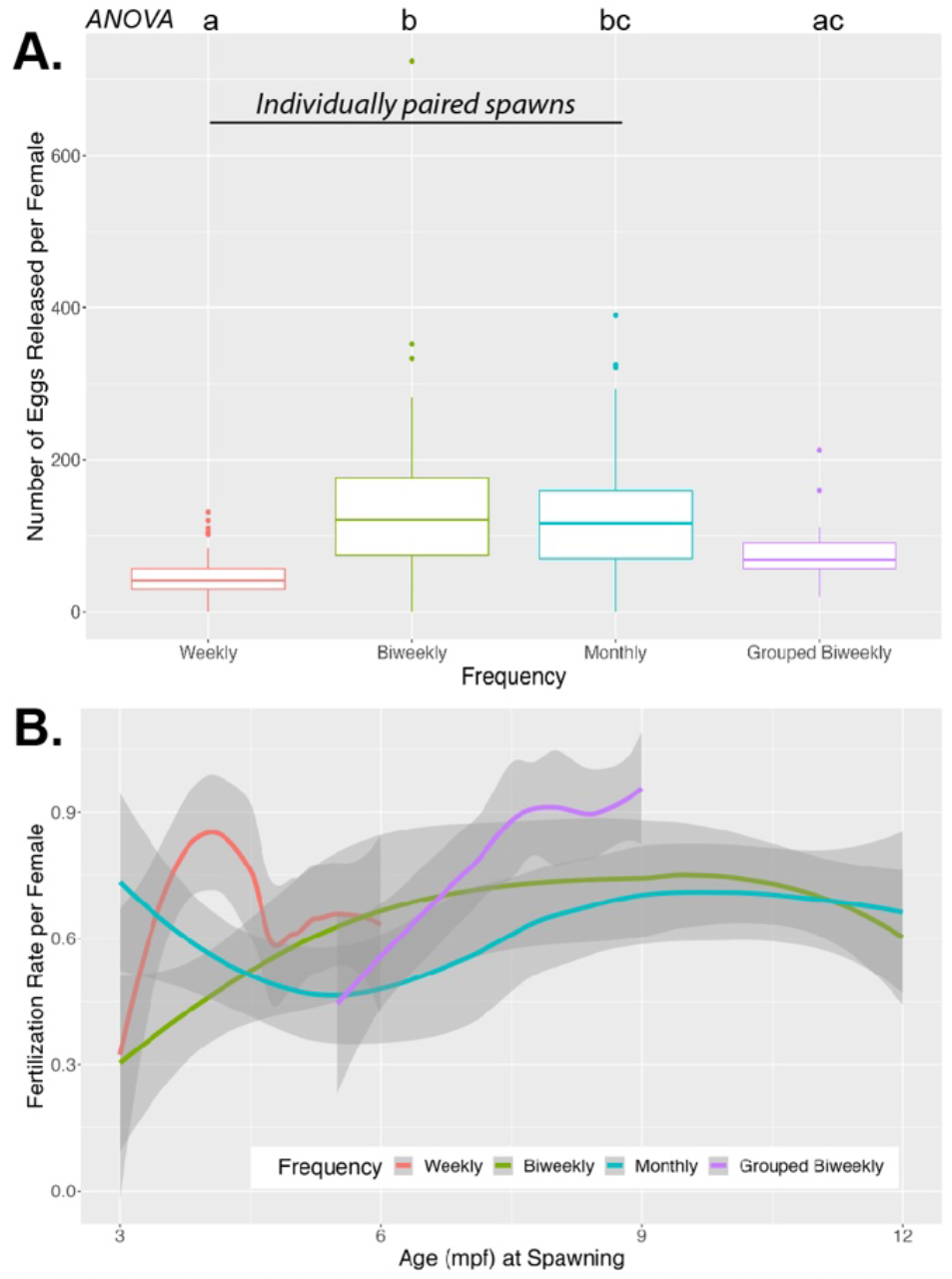
Spawning frequency significantly affects measures of female fertility. **(A)** The number of eggs released significantly varies with the frequency of spawning, but not at age spawning (one-way ANOVA, Tukey’s post-hoc analysis, p>0.05). **(B)** The fertilization rate is significantly affected by age and the frequency of spawning. Biweekly and monthly individually paired spawning have significantly different fertilization rates than weekly individually paired spawns or grouped biweekly spawns (linear regression, p<0.05). All spawnings were performed with multiple individuals and repeated with 2-3 different cohorts. The lines represent the mean and the gray bars are 95% confidence intervals.

The fertilization rate per female is another critical parameter used as an indicator of fertility. Age significantly affected the fertilization rate (p=0.00034, linear regression); therefore, the fertilization rate over the course of longitudinal experiments should not be combined. Compared to weekly individually paired spawns, biweekly and monthly individually paired spawns resulted in significantly different fertilization rates (p=0.00358, p=0.00591, respectively, linear regression, **Figure 1B**). This suggests that studies testing fertilization rates cannot be compared unless they are also using the same spawning frequency *and* age. Our literature search demonstrated that age, like all other factors of fertility testing, is inconsistent in the field.

Group spawns with equal sex ratios were also highly utilized in the literature. Based on the results from individually paired spawns, we hypothesized that the fertility measurements from group spawns would be affected by age and frequency, and would not be equivalent to measurements derived from individually paired spawns. We chose to compare biweekly individually paired spawns with biweekly grouped spawns as a representation of group vs. individually paired spawns. Surprisingly, the number of eggs released in biweekly grouped spawns was significantly different when compared to biweekly individually paired spawns (**Figure 1A**), but not different from weekly or monthly individually paired spawns. While the fertilization rate per female in biweekly grouped spawns was not different from weekly individually paired spawns (p=0.6529, linear regression). We did not test if varying the ratio of males to females also changed the number of eggs released, because this mating paradigm was not as widely used in the literature.

The age at which eggs are assessed for successful fertilization can potentially yield different information about the causes of reduced fertility. We assess fertility at approximately 4 hours post fertilization (hpf), in line with the procedures described in most studies (Table 1). However, other studies were not specific about when fertility assessment occurred, or, in some cases, the researchers waited until 3-5 dpf to assess the fertilization rate of a clutch. Fertilization occurs during spawning and the first few cell divisions can be observed within an hour of spawning. Activation of the zygotic genome occurs at 3 hpf and failure to develop past this stage can be considered teratogenicity [21,22]. Fertilization and early development are not interchangeable, therefore, the time point at which the eggs are evaluated will shape the conclusions that can be made about the effect of exposure on fertility.

In reviewing the literature, continuous spawning was often used to test fertility. Continuous spawning is not used in standard laboratory spawning practices, where it is generally understood that the females need approximately a week to regenerate eggs and there is often a need for timed matings for developmental studies. Continuous spawning for 21 days is recommended by the Organization for Economic Co-operation and Development (OECD) Guideline for the Testing of Chemicals, “Fish short-term reproduction assay” [23]. OECD recommends zebrafish are continuously spawned for 21 days at a ratio of 5 males to 5 females. Fish should be ∼4 mpf (+/-2 weeks) and fertilization should be assessed “Daily”. The US Environmental Protection Agency (USEPA) Endocrine Disruptor Screening Program guidelines suggest a 21-day continuous spawn (2 males to 4 females) of fathead minnow or medaka from ∼4.5-6 mpf for the short-term reproductive assay; however, the USEPA does not include zebrafish as a test species for this assay [24]. Further, the USEPA divides fertility into fecundity (the number of eggs per female), and fertilization rate (the number of embryos successfully fertilized divided by the total number of eggs), and fertilization can be observed from cleavage stage through 48 hpf. Although there are discrepancies between the OECD and USEPA guidelines, there are several points of consensus that can account for many of the studies in Table 1. The presence of these terms in the OECD and USEPA guidelines likely accounts for the use of this approach in many studies however the variable ages and time points recommended by the guidelines may not be comparable across studies.

Finally, the studies summarized in Table 1 also include inconsistent parameters that may interfere with the fertility assessment and comparison across studies. Inconsistencies included: additional spawning that was not recorded [5], randomly selected females and males so that a female may have been selected for consecutive spawns [4], and varying spawning conditions over time without a control group [1]. Our study showed that the frequency of spawning and age significantly altered measures of fertility; therefore, we recommend all fish used in fertility studies should be consistently spawned at a specific frequency without randomization or removing females if they do not spawn, and should be compared with an age-matched control.

Results for this study echo the chorus within the zebrafish toxicology research community that inconsistencies in experimental design have the potential to alter results and make it more difficult to compare studies from different labs. Studies have reported how rearing conditions have altered developmental timelines, behavioral studies, and toxicity assays [25]. Without an agreed-upon standard of care or methodology, these issues will continue to contribute to issues of rigor and reproducibility. In reference to studies of fertility, there are several distinct steps that can be taken to increase rigor and reproducibility. Manuscripts should include (1) the male-to-female ratio, (2) the frequency of spawning, (3) the specific time or window of time in which fertilization of eggs was determined, (4) the exact age spawning trails were initiated, and (5) the strain of zebrafish used, including the presence of any transgenes.

We found that the number of eggs released per female was the most consistent measure of fertility over time. However, the number of eggs released is not a measurement of fertilization rate or early development; therefore, the measure of fertility should be chosen based on the hypothesis being tested. In conclusion, zebrafish are a powerful tool in the study of fertility and endocrine disruption. By generating consensus for consistent practices in fertility testing, zebrafish have the potential to become a more influential model for assessing toxicant-induced changes in fertility.

## Data Availability

Data collected from the spawns performed in this study are available at: https://doi.org/10.26300/16gq-qc31.

## Funding Acknowledgement

MK was supported by F32ES023650 and 5T32ES007272 both from NIEHS. JP was supported by a CPVB Phase II COBRE (2PG20GM103652), and an NIEHS ONES award (ES030109).

